## supplemental Files for "A Novel Genus of Virulent Phage Targeting *Acinetobacter baumannii*: Efficacy and Safety in a Murine Model of Pulmonary Infection"

Original article

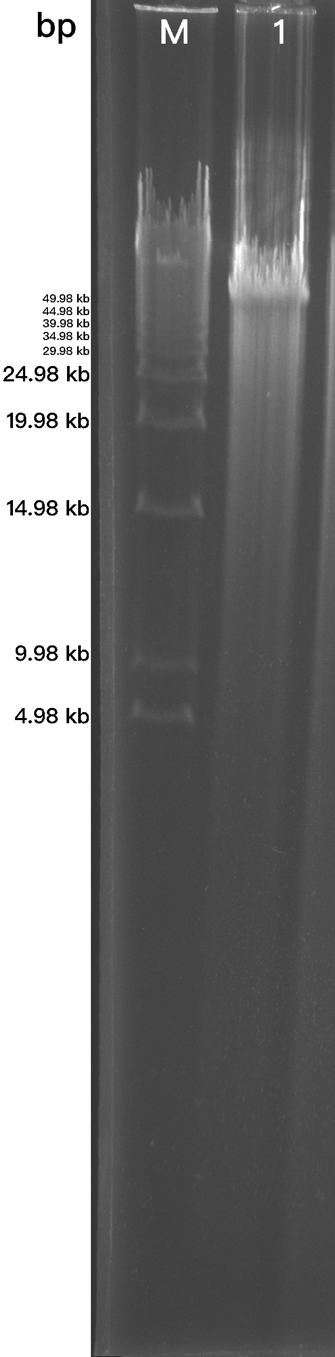

Figure S1: Pulsed-field electrophoresis of vB_AbaS_qsb1 genome.

M: DNA Size Standards - 5 kb ladder (BIO-RAD);
1: vB_AbaS_qsb1 genome.

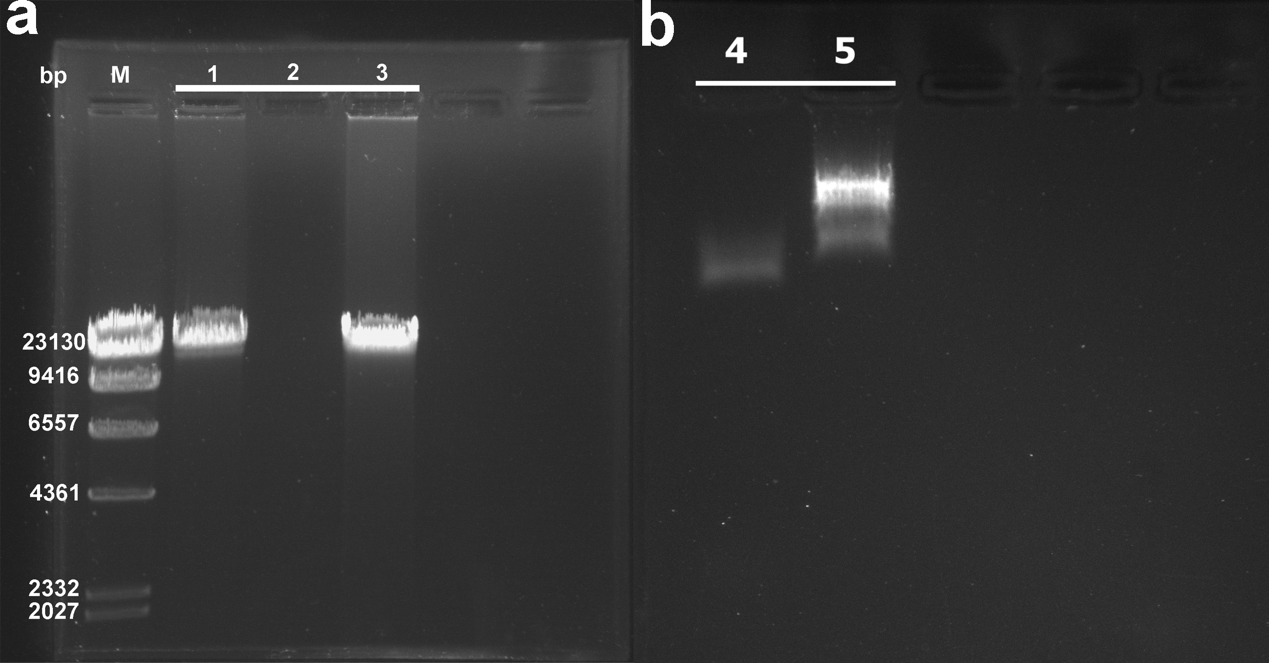

Figure S2: Validation of the nuclear type of vB_AbaS_qsb1;

1: vB_AbaS_qsb1 genome

2: DNAse I treatment

3: RNAse A treatment

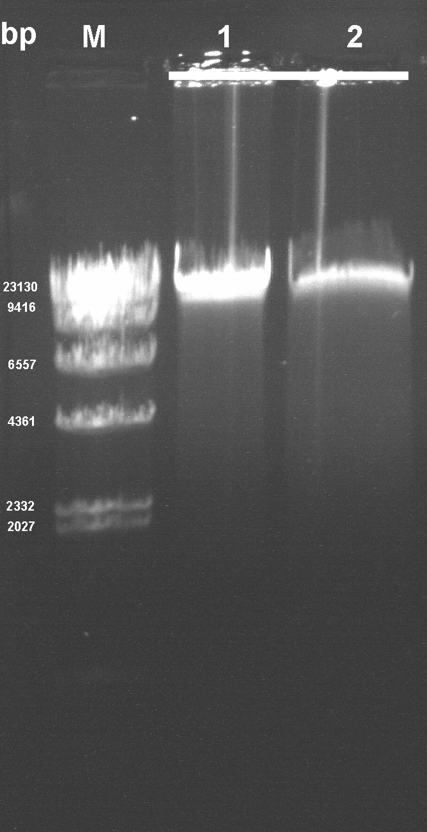

Figure S3: Validation of the nucleic acid structure of vB_AbaS_qsb1;

1: vB_AbaS_qsb1 genome

2: S1 nuclease treatment

Table S1: Genome-wide gene annotation information for vB_AbaS_qsb1

| name | Function classification | start | stop | strand | name |
| --- | --- | --- | --- | --- | --- |
| CDS_1 | integration | 169 | 1383 | + | Integrase Arm-type DNA-binding domain |
| CDS_2 | CDS | 1380 | 1571 | - | hypothetical protein |
| CDS_3 | CDS | 1707 | 2066 | + | hypothetical protein |
| CDS_4 | CDS | 2076 | 2507 | + | hypothetical protein |
| CDS_5 | CDS | 2504 | 2887 | + | hypothetical protein |
| CDS_6 | lysis | 3266 | 3820 | - | Phage Lysozyme RrrD |
| CDS_7 | CDS | 3817 | 4290 | - | hypothetical protein |
| CDS_8 | infection | 4274 | 4999 | - | putative side tail fiber protein |
| CDS_9 | CDS | 4999 | 5322 | - | hypothetical protein |
| CDS_10 | infection | 5323 | 9273 | - | Phage tail tip, host specificity protein J |
| CDS_11 | assembly | 9330 | 9983 | - | Phage tail tip, assembly protein I |
| CDS_12 | infection | 9967 | 10722 | - | tail assembly protein K |
| CDS_13 | infection | 10729 | 11535 | - | Phage tail tip, assembly protein L |
| CDS_14 | infection | 11522 | 12697 | - | Phage non-contractile tail fiber protein |
| CDS_15 | CDS | 12816 | 12962 | - | hypothetical protein |
| CDS_16 | Regulation | 13025 | 13348 | - | T3SS negative regulator,GrlR |
| CDS_17 | infection | 13424 | 13765 | - | Phage minor tail protein |
| CDS_18 | CDS | 13823 | 13942 | - | hypothetical protein |
| CDS_19 | CDS | 14178 | 14438 | - | hypothetical protein |
| CDS_20 | assembly | 14449 | 17226 | - | tail length tape measure protein |
| CDS_21 | CDS | 17335 | 17841 | - | hypothetical protein |
| CDS_22 | CDS | 17890 | 18714 | - | hypothetical protein |
| CDS_23 | CDS | 18707 | 19534 | - | hypothetical protein |
| CDS_24 | replication | 19570 | 19812 | - | Arc-like DNA binding domain protein |
| CDS_25 | replication | 19920 | 20075 | + | Arc-like DNA binding domain protein |
| CDS_26 | Regulation | 20146 | 21033 | + | BRO family, N-terminal domain protein |
| CDS_27 | CDS | 21137 | 21991 | - | hypothetical protein |
| CDS_28 | CDS | 22151 | 21954 | - | hypothetical protein |
| CDS_29 | assmbly | 22597 | 23091 | - | Tail assembly chaperone |
| CDS_30 | infection | 23107 | 24036 | - | putative major tail protein |
| CDS_31 | CDS | 24103 | 24348 | - | hypothetical protein |
| CDS_32 | replication | 24345 | 24887 | - | Phage antirepressor protein AntB |
| CDS_33 | infection | 25133 | 25531 | - | tail terminator protein |
| CDS_34 | assembly | 25533 | 25988 | - | tail completion or Neck1 protein |
| CDS_35 | CDS | 26027 | 26698 | - | hypothetical protein |
| CDS_36 | assembly | 26754 | 27134 | - | head-tail joining protein |
| CDS_37 | assembly | 27137 | 27529 | - | head-tail adaptor Ad1 |
| name | Function classification | start | stop | strand | name |
| CDS_38 | assembly | 27533 | 27946 | - | LEM-like domain protein |
| CDS_39 | assembly | 27986 | 29140 | - | major capsid protein |
| CDS_40 | assembly | 29153 | 29959 | - | minor structural protein GP20 |
| CDS_41 | CDS | 30067 | 30219 | - | hypothetical protein |
| CDS_42 | CDS | 30235 | 30465 | - | hypothetical protein |
| CDS_43 | assembly | 30462 | 31547 | - | putative head morphogenesis protein |
| CDS_44 | Regulation | 31513 | 32916 | - | portal protein |
| CDS_45 | packaging | 32924 | 34420 | - | terminase large subunit |
| CDS_46 | packaging | 34436 | 34897 | - | terminase small subunit |
| CDS_47 | CDS | 34947 | 35729 | - | hypothetical protein |
| CDS_48 | CDS | 35801 | 35923 | - | hypothetical protein |
| CDS_49 | CDS | 35951 | 36133 | - | hypothetical protein |
| CDS_50 | integration | 36188 | 36706 | - | NinB/ Orf homologous recombination mediator |
| CDS_51 | CDS | 36836 | 37300 | - | hypothetical protein |
| CDS_52 | packaging | 37355 | 37636 | - | HNH endonuclease |
| CDS_53 | CDS | 37587 | 37790 | - | hypothetical protein |
| CDS_54 | CDS | 37794 | 37982 | - | hypothetical protein |
| CDS_55 | tRNA | 38075 | 38162 | - | tRNA-Ile-TAT |
| CDS_56 | tRNA | 38168 | 38241 | - | tRNA-Arg-TCT |
| CDS_57 | CDS | 38294 | 38569 | - | hypothetical protein |
| CDS_58 | CDS | 38886 | 39440 | - | hypothetical protein |
| CDS_59 | CDS | 39451 | 39744 | - | hypothetical protein |
| CDS_60 | regulation | 39754 | 40179 | - | VRR-NUC domain protein |
| CDS_61 | regulation | 40183 | 40569 | - | HNH endonuclease |
| CDS_62 | CDS | 40569 | 40871 | - | hypothetical protein |
| CDS_63 | CDS | 40868 | 41002 | - | hypothetical protein |
| CDS_64 | regulation | 40999 | 41964 | - | putative DNA mismatch endonuclease |
| CDS_65 | CDS | 41961 | 42164 | - | hypothetical protein |
| CDS_66 | CDS | 42161 | 42562 | - | hypothetical protein |
| CDS_67 | CDS | 42564 | 42767 | - | hypothetical protein |
| CDS_68 | CDS | 42764 | 42970 | - | hypothetical protein |
| CDS_69 | CDS | 42963 | 43343 | - | hypothetical protein |
| CDS_70 | CDS | 43340 | 43510 | - | hypothetical protein |
| CDS_71 | CDS | 43507 | 43683 | - | hypothetical protein |
| CDS_72 | CDS | 43680 | 43937 | - | hypothetical protein |
| CDS_73 | replication | 43934 | 45259 | - | replicative DNA helicase |
| CDS_74 | replication | 45259 | 46152 | - | DnaD domain protein |
| name | Function classification | start | stop | strand | name |
| CDS_75 | CDS | 46149 | 46316 | - | hypothetical protein |
| CDS_76 | regulation | 46413 | 46679 | - | transcriptional regulator |
| CDS_77 | regulation | 46690 | 46920 | - | Regulatory protein Cro |
| CDS_78 | regulation | 47045 | 47791 | + | CI-like repressor |
| CDS_79 | CDS | 47805 | 48023 | + | hypothetical protein |
| CDS_80 | CDS | 48243 | 48434 | + | hypothetical protein |
| CDS_81 | CDS | 48434 | 48739 | + | hypothetical protein |
| CDS_82 | CDS | 48732 | 49454 | + | hypothetical protein |
| CDS_83 | CDS | 49451 | 49831 | + | hypothetical protein |
| CDS_84 | CDS | 49831 | 50028 | + | hypothetical protein |
| CDS_85 | CDS | 50025 | 50210 | + | hypothetical protein |
| CDS_86 | CDS | 50210 | 50539 | + | hypothetical protein |
| CDS_87 | CDS | 50549 | 51487 | + | hypothetical protein |
| CDS_88 | packaging | 51484 | 52143 | + | exonuclease |
| CDS_89 | CDS | 52140 | 52403 | + | hypothetical protein |
| CDS_90 | CDS | 52415 | 52780 | + | hypothetical protein |
| CDS_91 | CDS | 52777 | 52986 | + | hypothetical protein |
| CDS_92 | CDS | 52983 | 53174 | + | hypothetical protein |
| CDS_93 | regulation | 53175 | 53375 | + | Prophage regulatory protein |
| CDS_94 | regulation | 53477 | 53950 | + | SsrA-binding protein |
| CDS_95 | regulation | 53992 | 54543 | - | flavodoxin-like protein |

#### Table S2: Partial genome alignment results obtained by BLAST

| Contigs | Query length (bp) | Query cover |
| --- | --- | --- |
| TPA_asm: *Caudoviricetes* sp. isolate ctGTK9 | 12320 | 49% |
| *Acinetobacter* phage Aclw_8 | 47018 | 14% |
| *Acinetobacter* phage TCUAN1 | 49691 | 30% |
| TPA_asm: *Caudoviricetes* sp. isolate ctMhV2 | 49432 | 12% |
| *Acinetobacter* phage Acba_16 | 51595 | 9% |
| *Acinetobacter* phage Acba_1 | 50696 | 6% |
| *Acinetobacter* phage Ab105-3phi | 63785 | 2% |
| TPA_asm: *Caudoviricetes* sp. isolate ctTRl6 | 36165 | 2% |
| TPA_asm: *Caudoviricetes* sp. isolate ctCs82 | 47750 | 0% |
| TPA_asm: *Caudoviricetes* sp. isolate ct6H824 | 42126 | 0% |
| TPA_asm: *Siphoviridae* sp. ct2u94 | 49650 | 0% |
| TPA_asm: Bacteriophage sp. isolate ctcvu2 | 185987 | 0% |
| TPA_asm: *Caudoviricetes* sp. isolate ctSIK6 | 41285 | 0% |
| TPA_asm: *Caudoviricetes* sp. isolate ctyJT3 | 42806 | 0% |

Table S3：Mean AAI value of phage of *Vieuvirus* genus

| Contigs（genus of *Vieuvirus*） | Mean AAI value (%) |
| --- | --- |
| *Acinetobacter* phage Aclw_8 | 73.07 |
| *Acinetobacter* phage Ab11510-phi | 70.41 |
| *Acinetobacter* phage A3926.3 | 63.26 |
| *Acinetobacter* phage vB_AbaS_Eva | 63.37 |
| *Acinetobacter* phage vB_AbaS_Ftm | 61.19 |
| *Acinetobacter* phage vB_AbaS_SA1 | 63.44 |
| *Acinetobacter* bacteriophage YC#06 | 58.65 |
| *Acinetobacter* bacteriophage YMC/09/02/B1251_ABA_BP | 58.90 |
| *Acinetobacter* bacteriophage YMC11_11_R3177 | 70.74 |

Table S4: Mean AAI value of phage of *Obolenskvirus* genus

| Contigs（genus of *Obolenskvirus*） | Mean AAI value (%) |
| --- | --- |
| *Acinetobacter* phage BUCT628 | 32.15 |
| *Acinetobacter* phage AB1 | 33.41 |
| *Acinetobacter* phage AP22 | 62.75 |

#### Table S5：OrthoANIu Results against vB_AbaS_qsb1

| Contigs | Total length (bp) | GC content (%) | OrthoANIu value (%) |
| --- | --- | --- | --- |
| *Acinetobacter* bacteriophage YMC11_11_R3177 | 45,575 | 39.83 | 73.50 |
| *Acinetobacter* bacteriophage YMC/09/02/B1251_ABA_BP | 45,364 | 39.05 | -1 |
| *Acinetobacter* bacteriophage YC#06 | 48,885 | 40.14 | 60 |
| *Acinetobacter* bacteriophage AP22 | 46387 | 37.74 | 73.40 |
| *Acinetobacter* bacteriophage AB1 | 45159 | 37.69 | -1 |
| *Acinetobacter* phage BUCT628 | 44935 | 37.52 | -1 |

**Material and method**

**Determination of bacterial and phage colonization**

After the mice were sacrificed, their lungs were aseptically removed, weighed, and mechanically homogenized in PBS containing protease inhibitors (PMSF). The lung homogenates were kept on ice and serially diluted within 1 hour. The bacterial load (CFU) in the lungs was quantified using standard microbiological methods, while the phage titer (PFU) of bacteriophage qsb1 was determined using the DLA method. Finally, the unit is CFU/g or PFU/g.

**Cytokine quantification**

For the treated lung tissue homogenate, the concentrations of interleukins (IL-1β, IL-6) and tumor necrosis factor-α (TNF-α) were quantified using ELISA. Cytokine levels were expressed the total protein relative content in the lung homogenate, measured as the optical density at 450 nm (OD_450nm_).

**Complete blood count**

After collecting whole blood from the mice, blood parameters were analyzed within 1 hour using an automated blood cell counter (XN-550,Sysmex America).

**Histopathological Analysis**

Following euthanasia, lung tissues were harvested and immediately fixed in 4% paraformaldehyde. The fixed tissues were processed for dehydration using an automated tissue dehydration processor (HistoCore PEGASUS, Leica), embedded in paraffin (HistoCore Arcadia H&C, Leica), and sectioned into 4 μm thick slices using a microtome (HistoCore Multicut, Leica). Sections were stained with hematoxylin and eosin (H&E) utilizing an automated staining system (HistoCore CHROMAX ST, Leica) and coverslipped with an automated coverslipper (Leica CV5030). Digital images of the stained sections were acquired and analyzed using a digital pathology slide scanner (KF-PRO-005-EX, KFBIO).
